# Proteomic and biochemical comparison of the cellular interaction partners of human VPS33A and VPS33B

**DOI:** 10.1101/236695

**Authors:** Morag R. Hunter, Geoffrey G. Hesketh, Anne-Claude Gingras, Stephen C. Graham

## Abstract

Multi-subunit tethering complexes control membrane fusion events in eukaryotic cells. CORVET and HOPS are two such multi-subunit tethering complexes, both containing the Sec1/Munc18 protein subunit VPS33A. Metazoans additionally possess VPS33B, which has considerable sequence similarity to VPS33A but does not integrate into CORVET or HOPS complexes and instead stably interacts with VIPAR. It has been recently suggested that VPS33B and VIPAR comprise two subunits of a novel multi-subunit tethering complex (named ‘CHEVI’), analogous in configuration to CORVET and HOPS. We utilised the BioID proximity biotinylation assay to compare and contrast the interactomes of VPS33A and VPS33B. Overall, few proteins were identified as associating with both VPS33A and VPS33B, suggesting these proteins have distinct sub-cellular localisations. Consistent with previous reports, we observed that VPS33A was co-localised with many components of class III phosphatidylinositol 3-kinase (PI3KC3) complexes: PIK3C3, PIK3R4, NRBF2, UVRAG and RUBICON. Although in this assay VPS33A clearly co-localised with several subunits of CORVET and HOPS, no proteins with the canonical CORVET/HOPS domain architecture were found to co-localise with VPS33B. Instead, we identified two novel VPS33B-interacting proteins, VPS53 and CCDC22. CCDC22 co-immunoprecipitated with VPS33B and VIPAR in over-expression conditions and interacts directly with the VPS33B-VIPAR complex in vitro. However, CCDC22 does not appear to co-fractionate with VPS33B and VIPAR in gel filtration of human cell lysates. We also observed that the protein complex in HEK293T cells which contained VPS33B and VIPAR was considerably smaller than CORVET/HOPS, suggesting that, unlike VPS33A, VPS33B does not assemble into a large stable multi-subunit tethering complex.

## INTRODUCTION

Membrane trafficking in eukaryotic cells is tightly controlled by a range of proteins, including markers of membrane identity (such as small GTPases) and multi-subunit tethering complexes. Tethering complexes, containing Sec1/Munc18 protein subunits, are soluble cytoplasmic proteins that work together with SNAREs to fuse membranes (1). One such Sec1/Munc18 protein is Vps33, found in all eukaryotes as a subunit of class C core vacuole/endosome tethering (CORVET) and homotypic fusion and vacuole protein sorting (HOPS) complexes (2). The mammalian HOPS and CORVET complexes share four common components, VPS33A, VPS16, VPS18 and VPS11, and have two distinct components, VPS8 and TRAP-1 (CORVET), or VPS39 and VPS41 (HOPS) (3–5). CORVET is active on Rab5-positive membranes (early endosomes) (3,6) and HOPS on Rab7-positive membranes (late endosomes, autophagosomes, lysosomes) (7–9).

In metazoans there are two Vps33 homologs (10), with the functions of yeast Vps33 being carried out by VPS33A. Metazoan VPS33B shares 30% amino acid sequence identity with VPS33A, but cannot integrate into CORVET or HOPS complexes (4,8,11). Instead, VPS33B has been observed on Rab10- and Rab25-positive membranes, functioning in post-Golgi trafficking (12), and on Rab11A-positive membranes acting in an apical recycling pathway (13).

Mutations in human VPS33A and VPS33B produce distinct phenotypes. Recently, a small population of patients have been identified with a single mutation in VPS33A (R498W) and a mucopolysaccharidosis-like phenotype (14,15). Although the mechanism is as yet unclear, this mutation results in over-acidification of lysosomes and an inability to catabolize glycosaminoglycans. This phenotype is quite different to that of patients with VPS33B mutations. VPS33B forms a complex with VIPAR (also known as SPE-39, VIPAS39 or VPS16B), and mutations in either of these proteins can cause Arthrogryposis, Renal dysfunction, and Cholestasis (ARC) syndrome (13). ARC syndrome is a multi-system disorder, with some symptoms attributable to the known functions of VPS33B and VIPAR in apical recycling pathways (13), post-Golgi collagen processing (12), and α-granule formation in megakaryocytes (16,17). Further, mutations in VPS33B that affect its interactions with Rab proteins cause Autosomal recessive Keratoderma-Ichthyosis-Deafness (ARKID) syndrome (18). The differences in phenotype conferred by mutations in VPS33A and VPS33B confirm that these proteins act in distinct cellular pathways.

A recent review suggested that VPS33B and VIPAR were members of a multi-subunit membrane tethering complex with an analogous organisation to CORVET and HOPS (19). This hypothetical complex was called “class C Homologs in Endosome-Vesicle Interaction” (CHEVI). With only two known subunits (VPS33B and VIPAR), there would be four other currently-unidentified subunits in this proposed complex. Here, we have used a quantitative proteomic approach in an attempt to identify members of the proposed CHEVI complex, and to further investigate the differences between the cellular membranes with which VPS33A and VPS33B associate as part of their cognate multi-subunit tethering complexes. We identified known interactors of VPS33A and VPS33B, and novel VPS33B interactors, but did not find evidence for the existence of the CHEVI complex.

## RESULTS

It is known that VPS33A must interact with VPS16 in order to be recruited to HOPS and to localise correctly to target membranes (8,11,20). Mutation of two residues of VPS33A (Y438D and I441K) is sufficient to prevent the interaction between VPS33A and VPS16, and therefore inhibit HOPS function (8). We utilised this construct as a negative control for VPS33A localisation in our BioID assays. The VPS33B-VIPAR complex is likely to closely resemble the VPS33A/VPS16 interaction (12). We therefore docked a model of VPS33B generated using I-TASSER (21) onto the structure of human VPS33A in complex with VPS16 (11) to identify point mutations that may disrupt the VPS33B-VIPAR interaction, thus preventing correct VPS33B localisation within the cell (figure 1A).

**Figure 1.**
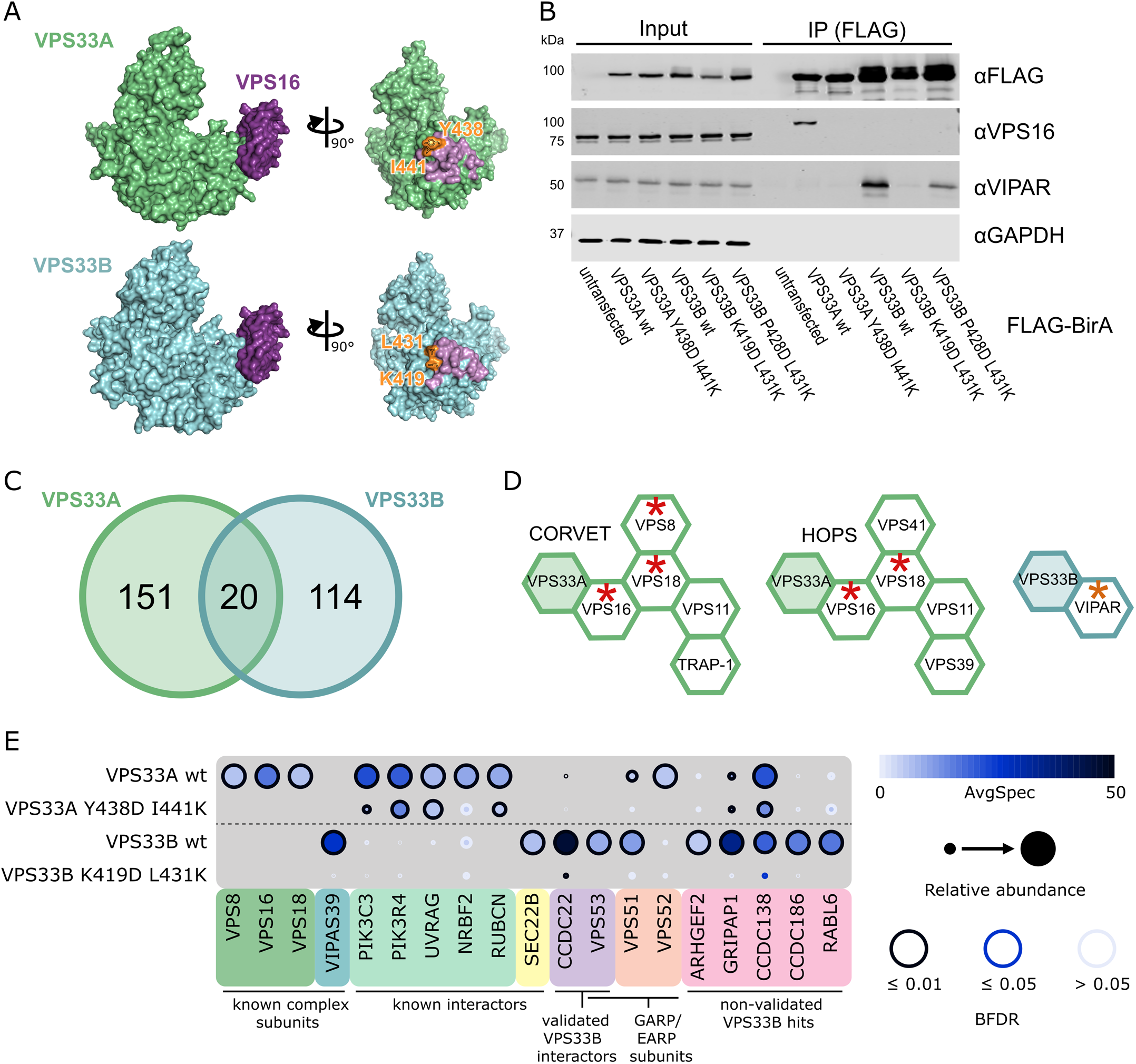
BioID results for VPS33A and VPS33B. (A) Crystal structure of VPS33A in complex with VPS16 residues 642–736 (top), and a homology model of VPS33B docked onto this complex (bottom). Residues shown in orange are mutated in figure B. Molecular graphics were generated using PyMOL (Schrodinger LLC). (B) HEK293 Flp-In T-REx cells were stably transfected with VPS33A and VPS33B (wildtype and mutant) constructs with C-terminal FLAG-BirA* tags and expression was induced with tetracycline. After immunoprecipitation (IP), samples were immunoblotted for the indicated endogenous protein. (C) Distribution of proteins identified by BioID with a Bayesian False Discovery Rate (BFDR) ≤0.01 and at least two-fold enrichment in the wildtype sample over the negative-control mutant sample. (D) CORVET, HOPS and VPS33B-VIPAR complexes, with subunits identified in BioID results marked with red asterisks (VPS33A as bait) or orange asterisks (VPS33B as bait). (E) Selected BioID results, shown as dot plots (55). The spectral counts for each indicated prey protein are shown as AvgSpec. VIPAS39 = VIPAR, RUBCN = RUBICON. A complete list of the proximal proteins for each bait is available in the supplementary data file.

In order to compare VPS33A and VPS33B, we utilised the proximity-based ligation (BioID) assay (22). This assay utilises an abortive biotin ligase, BirA*, which releases an active intermediate of biotin (biotinoyl-5’-AMP) that can covalently label primary amines on proteins in the vicinity of (but not necessarily in direct contact with) the tagged protein of interest. We anticipated that BirA*-tagged VPS33A and VPS33B would result in biotinylation of stable tethering complex subunits and markers of membrane identity.

Wildtype and mutant VPS33A and VPS33B were C-terminally tagged with FLAG-BirA*, and stably integrated into the tetracycline-inducible HEK293 Flp-In T-REx cell line. As expected, when expressed in this cell line, wildtype VPS33A-FLAG-BirA* was able to co-immunoprecipitate endogenous VPS16, while the VPS33A Y438D I441K mutant could not (figure 1B). Similarly, wildtype VPS33B-FLAG-BirA* was able to co-immunoprecipitate endogenous VIPAR (figure 1B). The VPS33B K419D L431K mutant entirely disrupted binding to endogenously-expressed VIPAR, while the VPS33B P428D L431K mutant retained some binding (figure 1B). We thus proceeded to use VPS33A Y438D I441K and VPS33B K419D L431K mutants as negative controls for their respective wildtype constructs, thereby facilitating the identification of interactions formed only when VPS33A and VPS33B were correctly assembled into their cognate multi-subunit tethering complexes: CORVET and HOPS for VPS33A, and with VIPAR (potentially as part of CHEVI) for VPS33B.

High-confidence proximity interactions (preys) in BioID assays were considered as having a Bayesian false discovery rate (BFDR) of ≤0.01 (as determined by SAINT analysis (23)) and at least two-fold enrichment of the protein in the wildtype compared to the negative control mutant. Of these high-confidence preys, very few proteins (20 of 285, 7%) were captured by both VPS33A and VPS33B (figure 1C). The majority of preys were found exclusively with either VPS33A or VPS33B, providing strong evidence that VPS33A and VPS33B occupy distinct subcellular locations.

Proteins identified for VPS33A included two of the common subunits of CORVET and HOPS, VPS16 and VPS18, demonstrating that the placement of the BirA* tag did allow for biotinylation of central complex subunits (figure 1D). We also identified VPS8, a subunit specifically found in CORVET. Most of the subunits of CORVET and HOPS (excluding VPS33A) follow a similar architecture: an N-terminal β-propeller, an α-solenoid, and (often) a C-terminal zinc-finger domain (2,24). Although the mass spectrometry (MS) results for VPS33B included VIPAR (gene name VIPAS39), we did not find any other proteins with a domain structure reminiscent of that of CORVET or HOPS subunits (a β-propeller followed by an α-solenoid) as high-confidence preys.

In our BioID results for VPS33A, we identified two out of three of the core subunits of class III phosphatidylinositol 3-kinase (PI3KC3) complexes - PIK3C3 (a.k.a. Vps34) and PIK3R4 (a.k.a. Vps15). PI3KC3 complexes are found on Rab5- (25) and Rab7-positive membranes (26), including early endosomes and late endosomes or autophagosomes (reviewed in (27)) where CORVET and HOPS function, respectively. We also identified NRBF2, a regulator of the PI3KC3-C1 complex (28–30). Furthermore, we identified UVRAG, which is a member of the PI3KC3-C2 complex and is a known interactor of HOPS during autophagosome-lysosome fusion (31). RUBICON (gene name RUBCN), an inhibitor of PI3KC3-C2 complexes, was also identified as a VPS33A prey. RUBICON reportedly binds directly to UVRAG and prevents UVRAG from binding to the HOPS complex (32,33). Our observation that VPS33A can capture RUBICON indicates that UVRAG may not be required for HOPS recruitment to these membranes.

For VPS33B, we observed one previously-known interacting protein as a high confidence prey – SEC22B. SEC22B is a SNARE protein with functions in delivering ER-resident proteins to phagosomes in dendritic cells (34) and in plasma membrane expansion (35). It has recently been described as interacting with VPS33B during the formation of α-granules in megakaryocytes (36).

As the BioID results for VPS33A included many interactors that have already been characterised, we focussed on validating our high confidence hits from VPS33B, most of which had not been described previously. Seven proteins were selected for further investigation, based on those appearing with high abundance in our MS results and having a known involvement in membrane trafficking processes. N-terminal myc-tags were added to the shortlisted proteins ARHGEF2, CCDC22, CCDC138, CCDC186, GRIPAP1, RABL6, and VPS53. These were each co-transfected with VPS33B-GFP and FLAG-VIPAR into HEK293T cells, and then a GFP immunoprecipitation was performed (supplementary figure 1). Two of the shortlisted proteins were found to co-immunoprecipitate with VPS33B-GFP under these conditions: myc-CCDC22 and myc-VPS53 (figure S1A). Further, over-expression and co-immunoprecipitation of either of these proteins did not affect the co-immunoprecipitation of FLAG-VIPAR, suggesting that this interaction with VPS33B was not competing with VIPAR binding.

CCDC22 is a subunit of the CCC complex (also called the ‘Commander’ complex), along with CCDC93 and COMMD proteins (37,38). The CCC complex has been found to interact with other multi-subunit complexes: WASH, Retromer and Retriever (37,39-41). VPS53 is a subunit of both the Golgi associated retrograde protein (GARP) and endosome-associated recycling protein (EARP) complexes. Both GARP and EARP have four known subunits, sharing three common subunits (VPS51, VPS52 and VPS53) and each containing one unique subunit (VPS54 in GARP, VPS50 in EARP) (42). Interestingly, VPS51 was also a lower-confidence result in our BioID dataset for VPS33B, and VPS52 was a lower-confidence result for VPS33A.

To probe whether GFP-tagged VPS33B was competent to bind endogenous CCDC22 or VPS53, an immunoprecipitation was performed using only transfected VPS33B-GFP (figure 2). Endogenous CCDC22 was efficiently co-immunoprecipitated by VPS33B-GFP. The endogenous CCC complex proteins CCDC93 and COMMD1 were not efficiently captured, nor was the WASH complex protein FAM21. Additionally, endogenous VPS53 did not appear to co-immunoprecipitate with VPS33B-GFP. However, it should be noted that the antibody against endogenous VPS53 indicates that these HEK293T cells expressed VPS53 isoform 4 (94 kDa), while our over-expression constructs are isoform 1 (80 kDa).

**Figure 2.**
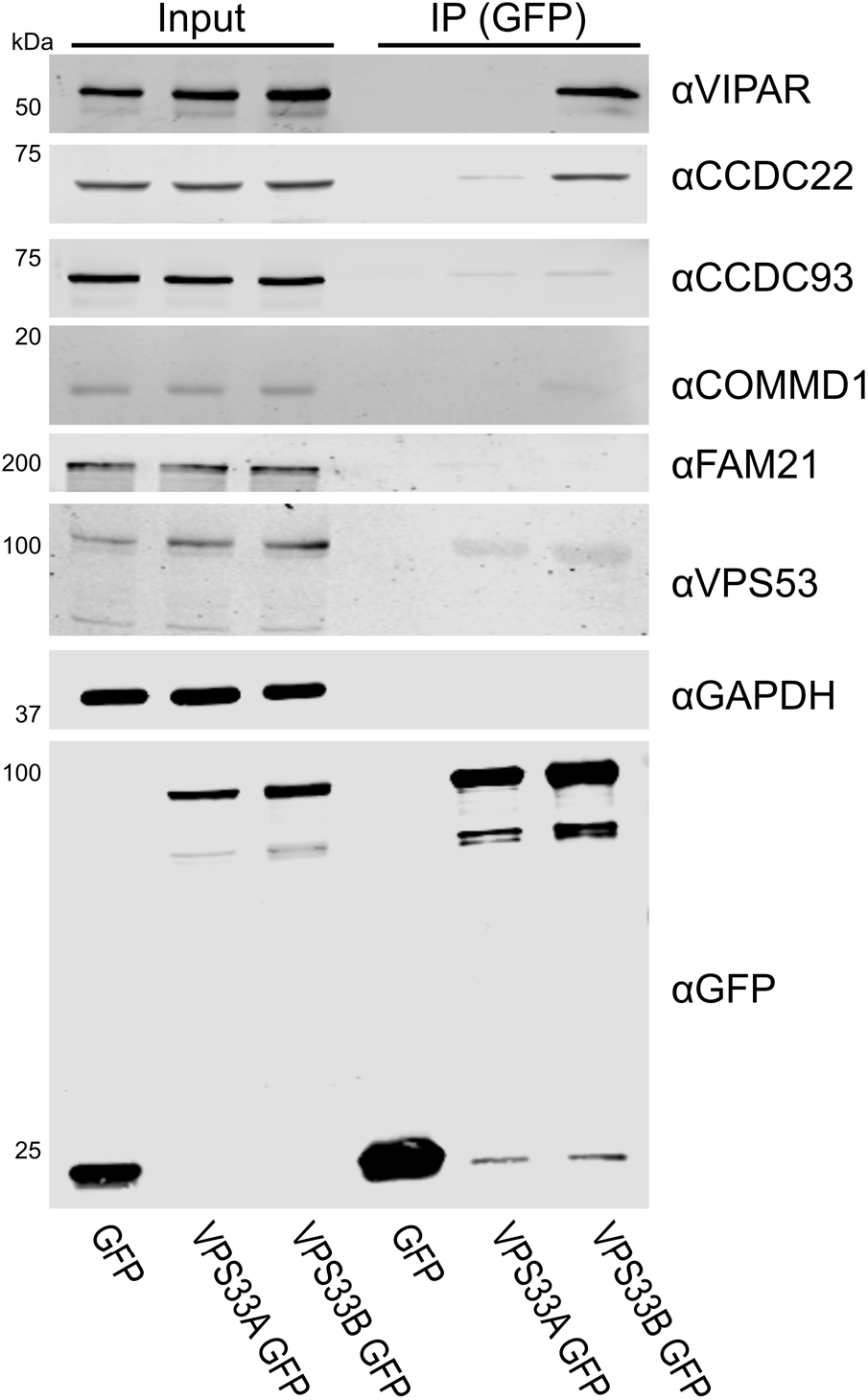
Endogenous CCDC22 is co-immunoprecipitated by VPS33B-GFP. (A) HEK293T cells were transfected with VPS33A-GFP or VPS33B-GFP. After immunoprecipitation (IP), samples were immunoblotted for endogenous proteins using the antibodies indicated.

In order to perform *in vitro* binding experiments, VPS33B and GST-VIPAR were co-expressed in *Escherichia coli* and purified by GSH affinity and size-exclusion chromatography (supplementary figure 2). The VPS33B/GST-VIPAR complex was used as bait in pulldown experiments, with myc-CCDC22 and myc-VPS53 expressed by cell-free in vitro transcription/translation in wheat germ lysate. Myc-CCDC22 was very efficiently pulled down by VPS33B/GST-VIPAR, whereas myc-VPS53 was not pulled down (figure 3A). This suggests that the interaction between VPS53 and the VPS33B-VIPAR complex is either indirect (i.e. other proteins contribute to the interaction) or requires a post-translational modification not conferred in the plant cell-free expression system. To test the former, we repeated the pulldown experiment with each of the other subunits of the GARP and EARP complexes (VPS50, VPS51, VPS52, VPS54), each with an N-terminal myc tag. However, none of these singly-expressed subunits interacted with VPS33B/GST-VIPAR (figure 3A). We also attempted co-expressing all the subunits of GARP or EARP simultaneously, in the hope of reconstituting the complexes, but this expression strategy was inefficient and did not result in binding to VPS33B/GST-VIPAR (data not shown).

**Figure 3.**
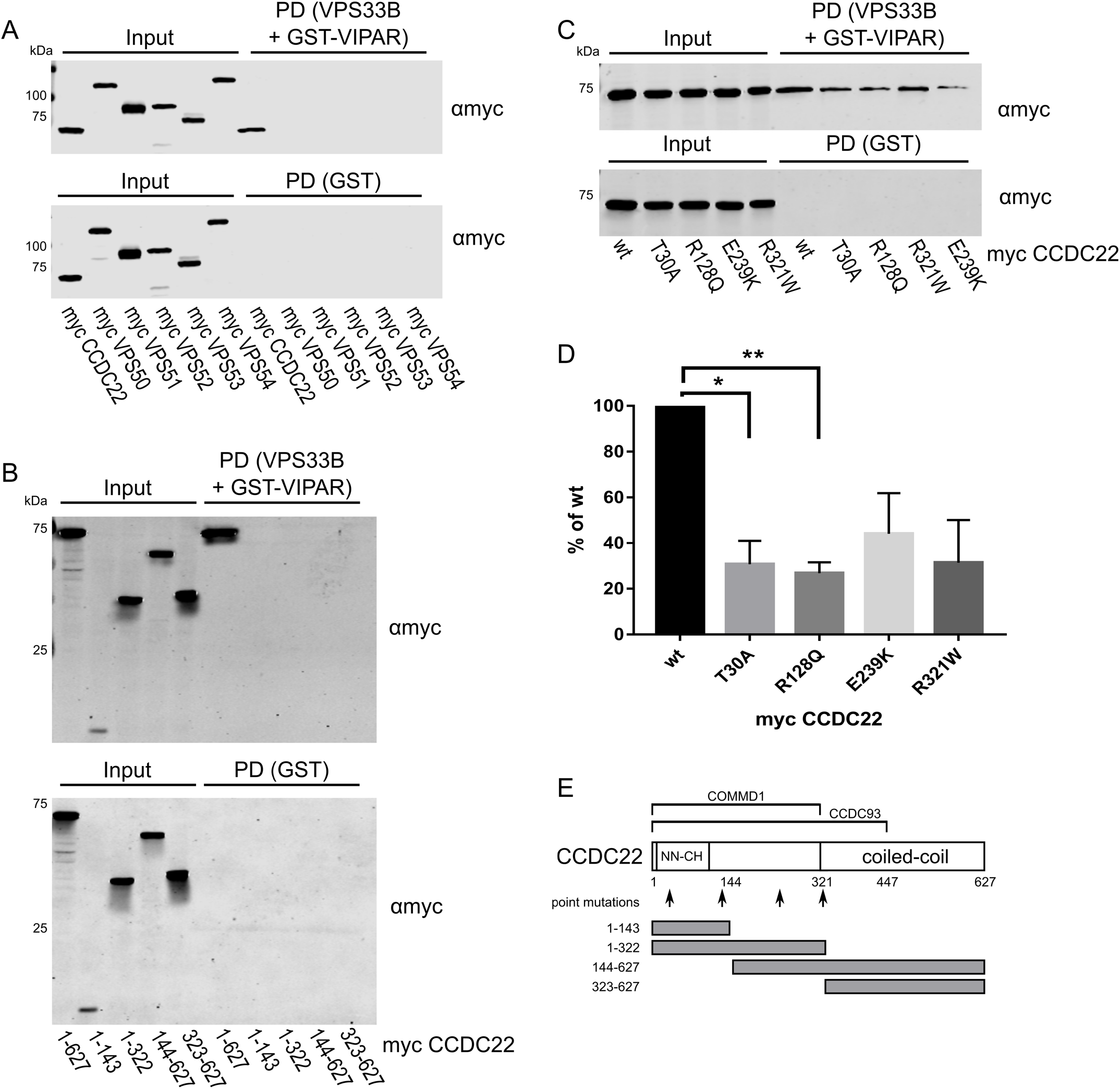
CCDC22 binds directly to VPS33B/GST-VIPAR in vitro. Myc-tagged CCDC22 or GARP and EARP subunits were produced by in vitro transcription/translation and then subjected to GST pulldown (PD) using VPS33B/GST-VIPAR or GST alone. Samples were analysed by immunoblotting with anti-myc. (A) CCDC22 is efficiently pulled down by VPS33B/GST-VIPAR, but no GARP or EARP subunits are pulled down. (B) While full length (1–627) CCDC22 is pulled down by VPS33B/GST-VIPAR, no CCDC22 truncations retained binding. (C) Point mutations of CCDC22 demonstrate reduced binding to VPS33B/GST-VIPAR compared to the wildtype (wt) protein. (D) Immunoblot band intensities were quantified and normalised (intensity of PD band divided by input band, normalised to the PD efficiency of the wildtype construct in each of three independent experiments), and analysed by one-way ANOVA with Dunnett’s multiple comparisons test (*p<0.05, **p<0.01). (E) Schematic of CCDC22, showing predicted N-terminal calponin homology (CH)-like (NN-CH) and C-terminal coiled-coil domains (59), regions required for binding to COMMD1 and CCDC93 (37), and constructs used in this paper.

To further understand how VPS33B-VIPAR may interact with CCDC22, we attempted to refine the region of CCDC22 that interacts with VPS33B/GST-VIPAR by generating a series of truncated forms of CCDC22. However, none of these CCDC22 truncations were able to bind to VPS33B/GST-VIPAR. We hypothesise that truncated forms of CCDC22 are unstable and unable to fold correctly in this assay system.

Multiple mutations have been reported in CCDC22, found in a cohort of X-linked intellectual disability (XLID) patients (43). Several of these mutations have been previously used to study the interaction between CCDC22 and COMMD proteins (found in the CCC complex) (44). Interestingly these mutations cluster in the N-terminal half of CCDC22. Four of these CCDC22 point mutants were assayed in the pulldown experiment described above, and each demonstrated reduced binding to VPS33B/GST-VIPAR. (The CCDC22 T17A mutant was not included because it also affects splicing (43).) For the two most N-terminal CCDC22 mutants, T30A and R128Q, we found statistically-significant disruption of binding to VPS33B/GST-VIPAR. This might suggest that the N terminus of CCDC22 is critical for binding to VPS33B/GST-VIPAR. However we cannot discount the possibility that these point mutations cause CCDC22 to be poorly folded in our assay system.

In order to determine whether VPS33B and VIPAR form a stable complex with CCDC22 in untransfected cells, we performed a cell fractionation experiment. In brief, whole cell lysates of HEK293T cells were separated by size exclusion chromatography. A series of elution fractions were collected from the column, the protein content of alternate fractions was concentrated, and the concentrated fractions were analysed by immunoblotting. In order from largest to smallest, we observed elution peaks for WASH, CORVET/HOPS, CCC, and then VPS33B-VIPAR complexes during cell lysate fractionation (figure 3A, 3B). It is thus evident that VPS33B and VIPAR do not co-fractionate with the majority of CCDC22 under these conditions. CCDC22 instead elutes in the same fractions as other members of the CCC complex (CCDC93 and COMMD1), proteins that are not efficiently co-immunoprecipitated by over-expressed VPS33B-GFP (figure 2). Further, the complex containing VPS33B and VIPAR elutes later than (and is thus considerably smaller than) the CORVET and HOPS complexes, indicating that VPS33B and VIPAR did not elute from the gel filtration column as part of a larger multi-subunit tethering complex.

## DISCUSSION

In this study we performed a proteomic comparison of the cellular interaction partners of VPS33A and VPS33B using proximity-based ligation (BioID). VPS33A was found to associate with members of the PI3K3C complexes, confirming previous observations (31–33). We also corroborated a recent publication that found VPS33B to co-localise with SEC22B (36). This confirms that our BioID-tagged constructs were able to localise to the correct intracellular membranes and were competent to label *bona fide* endogenous binding partners. Overall, the majority of proteins identified in our assay co-localised exclusively with either VPS33A or VPS33B, indicating that these proteins are found on different subcellular membranes.

VPS33A is a subunit of both the CORVET and HOPS complexes, the subunits of which share a very similar architecture (2,24). While VPS33B-FLAG-BirA* did label VIPAR, we did not find any other proteins proximal to VPS33B that shared the canonical architecture of CORVET and HOPS components, as would have been predicted in a VPS33B- and VIPAR-containing CHEVI complex. Further, our results from the whole cell fractionation experiment provide compelling evidence that VPS33B and VIPAR are found in a complex which is considerably smaller than CORVET and HOPS (figure 4). Given that CORVET/HOPS components co-eluted during this experiment, confirming that such complexes remain assembled under these assay conditions, we conclude that the hypothesised large, multi-subunit CHEVI complex does not exist in HEK293T cells.

**Figure 4.**
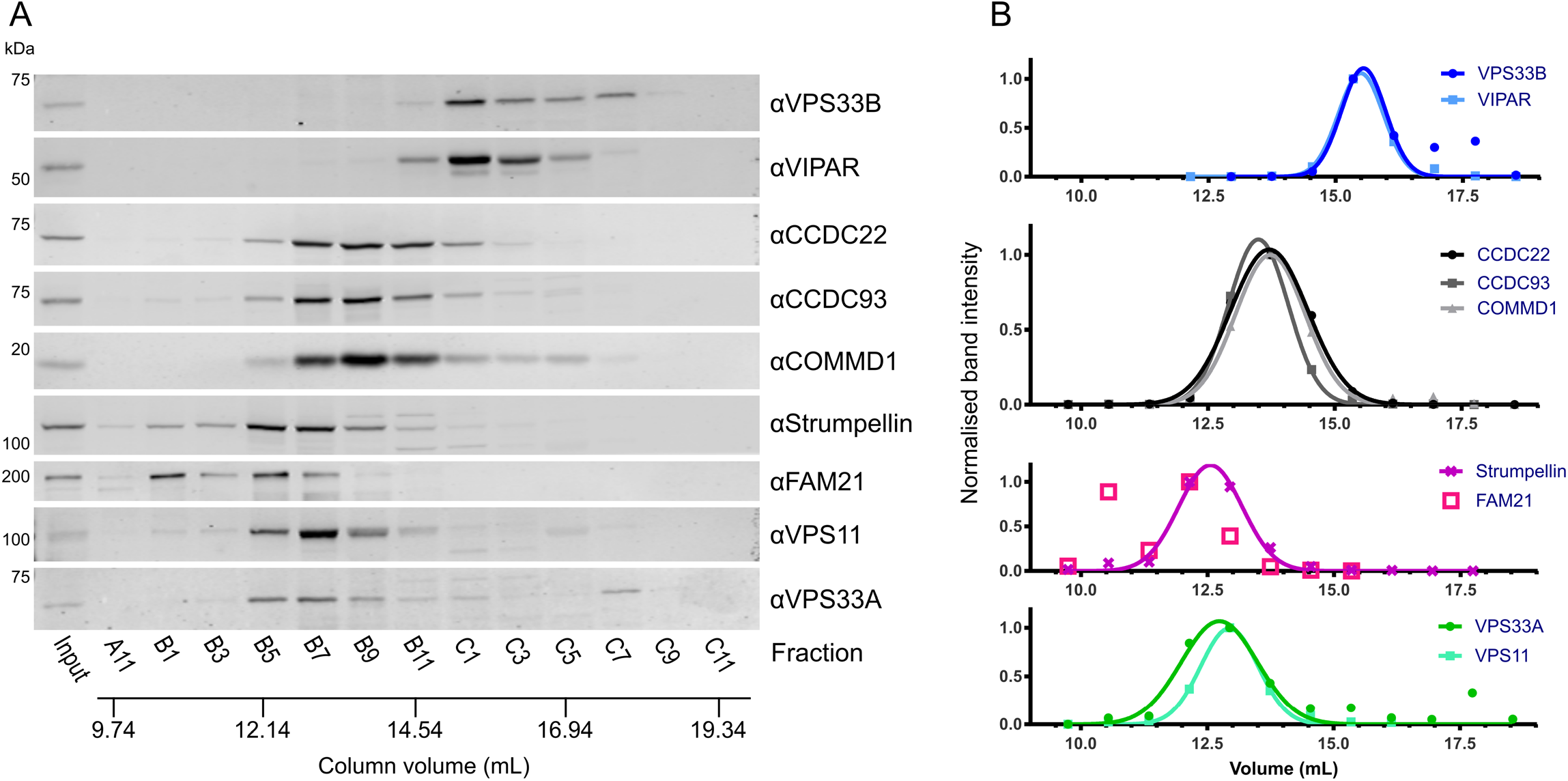
Whole cell fractionation of HEK293T cells shows that VPS33B and VIPAR form a complex which is considerably smaller than CORVET/HOPS and does not contain CCDC22. (A) HEK293T cell lysates were injected onto a Superose 6 10/300 GL gel filtration column and eluted fractions were analysed by immunoblotting. (B) Immunoblot band intensities were quantified, normalised to the band of maximum intensity, and fitted to a Gaussian distribution (for all except FAM21, which could not be reliably fitted to a single Gaussian distribution).

VPS33B and VIPAR have previously been described as acting in recycling pathways (12,13), as has VPS53 as part of GARP and EARP complexes (42,45), and CCDC22 as part of the CCC complex (37,39-41). Both VPS53 and CCDC22 associated with VPS33B and VIPAR when co-overexpressed (supplementary figure 1). Endogenous VPS53 was not co-immunoprecipitated by overexpressed VPS33B-GFP (figure 2), nor could the interaction with the VPS33B-VIPAR complex be observed *in vitro* (figure 3A), suggesting that this interaction is either transient or is tightly regulated by the cell. GFP-tagged VPS33B was able to co-immunoprecipitate endogenous CCDC22 and this interaction could be recapitulated using purified recombinant components (figures 2 and 3A), suggesting that it is a direct physical interaction. Mutations in CCDC22 that cause XLID reduce the interaction of this protein with the VPS33B-VIPAR complex (figure 3C–D). In agreement with our results, a recent large-scale proteomic study also identified CCDC22 as interacting with VPS33B-VIPAR (46). We do not observe co-immunoprecipitation of VPS33B-GFP with other members of the CCC complex (figure 2). Further, we do not observe efficient co-fractionation of endogenous VPS33B-VIPAR with CCDC22 from HEK293T cell lysates, with CCDC22 instead co-fractionating with other CCC complex components (figure 4). Notably, the CCC complex also did not co-fractionate with the WASH complex, a known interactor. This suggests that the interaction between VPS33B-VIPAR and CCDC22, either alone or as part of the CCC complex, may be weak or transient in cultured HEK293T cells. We hypothesise that this interaction may be inducible upon addition of an external stimulus, for example during bacterial infection, when VPS33B/VIPAR activity has been demonstrated to be important for a robust immunological response (47–49).

In summary, we propose that VPS33B-VIPAR is not assembled into a large multi-subunit tethering complex. Rather, VPS33B and VIPAR form a discrete bimolecular complex that can transiently interact with members of other multi-subunit trafficking complexes such as the CCC complex component CCDC22.

## EXPERIMENTAL PROCEDURES

### Expression constructs/cloning

Human VPS33A and VPS33B were cloned into pDONR223 (Invitrogen) and then pDEST-pcDNA5-BirA-FLAG (50) using Gateway cloning. VPS33A Y438D I441K, VPS33B K419D L431K and VPS33B P428D L431K were created by site-directed mutagenesis of wildtype constructs in pDONR223, then transferred into pDEST-pcDNA5-BirA-FLAG.

For transient transfection, wildtype VPS33A and VPS33B were cloned into pEGFP-N1 (Clontech), in order to add a C-terminal EGFP tag. An N-terminal FLAG-tag was added to VIPAR and cloned into pF5K CMV-neo (Promega). ARHGEF2 (isoform 1), CCDC22, CCDC138 (isoform 1), CCDC186, GRIPAP1 (isoform 1), RABL6 (isoform 1), and VPS53 (isoform 1) were cloned with N-terminal myc tags into pF5K CMV-neo.

For co-expression in *E. coli*, DNA encoding VPS33B and VIPAR that had been codon-optimised for bacterial expression (GeneArt) was cloned into positions 1 and 2 (respectively) into the polycistronic vector pOPC (51), VIPAR being tagged at the N terminus with GST.

For in vitro transcription/translation, full-length human CCDC22, VPS50, VPS51, VPS52, VPS53 and VPS54 were cloned into pF3A WG (BYDV) (Promega) with an N-terminal myc tag. Truncation and point mutations of CCDC22 were created by inverse PCR or site-directed mutagenesis.

### Cell culture and transfection

Untransfected HEK293 Flp-In T-REx cells were grown in Dulbecco’s Modified Eagle Medium with high glucose (Sigma, cat. D6546), supplemented with 10% (v/v) heat-inactivated foetal calf serum, 2 mM L-glutamine, 100 IU/ml penicillin, 100 mg/ml streptomycin, 3 μg/ml blasticidin and 100 μg/ml zeocin, in a humidified 5% CO2 atmosphere at 37°C. Cells were co-transfected with pOG44 (encoding the Flp recombinase) and the VPS33A or VPS33B constructs in pDEST-pcDNA5-BirA-FLAG vectors (described above) using TransIT-LT1 (Mirus), following the manufacturer’s instructions. Stable cell lines were selected by addition of 200 μg/ml hygromycin.

HEK293T cells (ATCC #CRL-3216, mycoplasma-free) were grown in DMEM with high glucose, supplemented with 10% (v/v) heat-inactivated foetal calf serum, 2 mM L-glutamine, 100 IU/ml penicillin and 100 mg/ml streptomycin, in a humidified 5% CO2 atmosphere at 37°C. Cells were transfected with TransIT-LT1 (Mirus), following the manufacturer’s instructions.

### BioID and mass spectrometry analysis

BioID and mass spectrometry analysis was performed essentially as described (52). Briefly, stable HEK293 Flp-In T-REx cells were grown on 15 cm plates to approximately 75% confluency. Bait expression and proximity labelling was then induced simultaneously by addition of tetracycline (1 μg/mL) and biotin (50 μM) for 24 hours. Cells were collected in PBS and biotinylated proteins were purified by streptavidin-agarose affinity purification. Proteins were digested on-bead with sequencing-grade trypsin in 50 mM ammonium bicarbonate pH 8.5. Peptides were then acidified by the addition of formic acid (2% (v/v) final concentration) and dried by vacuum centrifugation. Dried peptides were suspended in 5% (v/v) formic acid and analysed on a TripleTOF 5600 mass spectrometer (SCIEX) in-line with a nanoflow electrospray ion source and nano-HPLC system. Raw data were searched and analysed within ProHits LIMS (53) and peptides matched to genes to determine prey spectral counts (54). High confidence proximity interactions (BFDR ≤ 0.1) were determined through SAINT analysis (23) implemented within ProHits. Bait samples (biological duplicates) were compared against 12 independent negative control samples (6 BirA-FLAG only and 6 3xFLAG only expressing cell lines). The specific control samples used in this study were previously published as part of Chapat et al. (52). Dotplots were prepared in ProHits-Viz (55). Mass spectrometry data have been deposited in the MassIVE database (ID MSV000081814; available for FTP download at ftp://MSV000081814@massive.ucsd.edu) and with the ProteomeXchange Consortium (identifier PXD008457) (56). A summary of the results are attached as supplemental data.

### Recombinant protein expression and purification

In vitro protein expression was performed using TNT SP6 High Yield Wheat Germ reaction mix (Promega) as per the manufacturer’s instructions.

VPS33B and GST-VIPAR were co-expressed in *E. coli* B834(DE3). Bacteria were grown in 2x TY medium to an A600 of 0.8–0.9 at 37°C, cooled to 22°C, and protein expression was induced by the addition of 0.2 mM isopropyl β-d-thiogalactopyranoside. After 16–18 hr, cells were harvested by centrifugation at 5000 ×g for 15 min and the pellet was stored at −20°C until required.

Cells were thawed and resuspended in 20 mM Tris (pH 7.5), 300 mM NaCl, 0.5 mM MgCl_2_, 1.4 mM β-mercaptoethanol, 0.05% TWEEN 20, supplemented with 400 units of bovine DNase I (Sigma–Aldrich) and 200 μl of EDTA-free protease inhibitor mixture (Sigma–Aldrich) per 4–8 L of cell culture. Cells were lysed at 24 kpsi using a TS series cell disruptor (Constant Systems) and lysates were cleared by centrifugation at 40 000 ×g for 30 min at 4°C. Cleared lysate was incubated with glutathione sepharose 4B (GE Healthcare) for 1 h at 4°C, the beads were washed with 20 mM Tris (pH 7.5), 300 mM NaCl, and 1 mM DTT, and bound protein was eluted in wash buffer supplemented with 25 mM reduced glutathione. Protein was injected onto a Superdex 200 10/300 GL column (GE Healthcare) equilibrated in 20 mM Tris (pH 7.5), 200 mM NaCl, and 1 mM DTT. Fractions containing purified proteins were concentrated using 30 kDa nominal molecular mass cut-off centrifugal filter units (Millipore), diluted in glycerol to a final concentration of 50% (v/v) glycerol, and stored at −20°C until required. Purified protein complex identity was verified by peptide mass fingerprinting.

### Co-immunoprecipitations and pull-downs

FLAG immunoprecipitations were performed by harvesting cells and lysing in 10 mM Tris-HCl pH 7.5, 150 mM NaCl, 0.5 mM EDTA, 0.5% NP-40, and EDTA-free protease inhibitor cocktail (Sigma). Protein concentration in the resulting lysates was quantified by BCA assay (Thermo Scientific), and protein concentration was equalised before immunoprecipitation with anti-FLAG M2 magnetic resin (Sigma). 1 ml of whole cell lysate, containing approximately 1.9 mg of protein, was incubated with 30 μl of anti-FLAG resin for 1 hr at 4°C. Samples were then washed three times in wash buffer (10 mM Tris-HCl pH 7.5, 150 mM NaCl, 0.5 mM EDTA, 0.5% NP-40), then eluted by incubation with 250 μg/ml FLAG peptide (Sigma, cat. F3290).

GFP immunoprecipitations were performed using GFP-TRAP resin (ChromoTek), as in (57).

GST pulldown experiments were performed as described in (57), using 40-100 pmol purified VPS33B/GST-VIPAR complex as bait (described above).

### Antibodies and immunoblotting

The following primary antibodies were used for immunoblotting: polyclonal anti-GFP (Sigma, cat. G1544), monoclonal anti-FLAG M2 (Sigma, cat. 080M6035), monoclonal anti-myc (Millipore, cat. 05-724), monoclonal anti-GAPDH (Life Tech, cat. AM4300), polyclonal anti-VPS33A (as described in (11)), monoclonal anti-VPS33B (Santa Cruz, cat SC-398322), monoclonal anti-VIPAR (a.k.a. anti-SPE39, gift from S.W. L’Hernault, (58)), polyclonal anti-VPS16 (Santa Cruz, cat. sc-86939, lot I1913), polyclonal anti-CCDC22 (Proteintech, cat. 16636-1-AP), polyclonal anti-CCDC93 (Proteintech, cat. 20861-1-AP), polyclonal anti-COMMD1 (gift from E. Burstein, as described in (37)), polyclonal anti-FAM21 (Santa Cruz), polyclonal anti-Strumpellin (Santa Cruz), and polyclonal anti-VPS53 (Sigma cat. HPA024446, lot A106517). IRDye 800CW-conjugated secondary antibodies were supplied by LI-COR: goat anti-mouse (cat. 925-32210), goat anti-rabbit (cat. 925-32211), and donkey anti-rabbit (cat. 925-32213).

All samples were immunoblotted as per (57). Immunoblot band intensity was quantified using Image Studio Lite (version 5.4, LI-COR) and non-linear fitting to a Gaussian distribution was performed using Prism 7 (GraphPad).

### Whole cell fractionation

HEK293T cells were harvested and lysed in a buffer similar to that used for immunoprecipitation experiments: 10 mM Tris pH 7.5, 150 mM NaCl, 0.5 mM EDTA, 0.5% NP-40, EDTA-free protease inhibitor cocktail and Benzonase (Sigma). After centrifugation at 20 000 ×g for 10 min, lysates were injected onto a Superose 6 10/300 column (GE Healthcare) equilibrated in 10 mM Tris pH 7.5, 150 mM NaCl, 0.5 mM EDTA, 0.5% NP-40. Fractions (0.4 mL) were collected and concentrated using 10 μL of StrataClean resin (Agilent) per 100 μL of fractionated protein. Bound protein was eluted by boiling in SDS-PAGE loading buffer and samples were analysed by SDS-PAGE and immunoblotting.

## ACKNOWLEDGEMENTS

We thank S.W. L’Hernault, E. Burstein and M. Seaman for kindly providing reagents. This work was supported by a Sir Henry Dale Fellowship, jointly funded by the Royal Society and the Wellcome Trust, to SCG [098406/Z/12/Z] and an Isaac Newton Trust/Wellcome Trust ISSF/University of Cambridge Joint Research Grant to SCG. GGH was supported by a Parkinson Canada Basic Research Fellowship. ACG is a Tier 1 Canada Research Chair and supported by a CIHR Foundation grant (FDN 143301).

---

**Supplementary figure 1.**
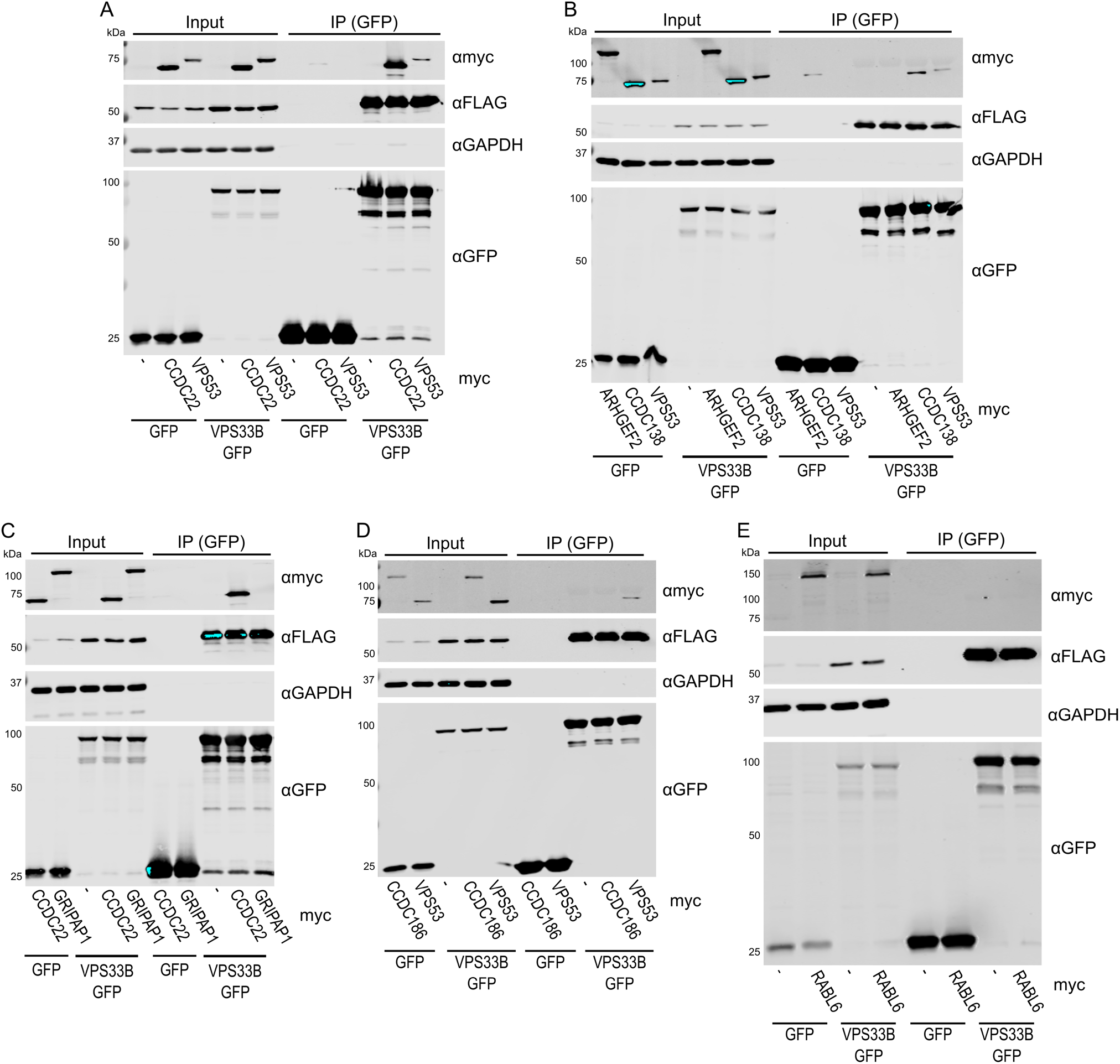
Validation of short-listed candidate VPS33B-interacting proteins. (A-E) ARHGEF2, CCDC22, CCDC138, CCDC186, GRIPAP1, RABL6, and VPS53 (with N-terminal myc tags) were each co-transfected with VPS33B-GFP and FLAG-VIPAR into HEK293T cells. A GFP immunoprecipitation (IP) was performed and samples were analysed by immunoblotting. Note that RABL6 appears above the expected size (81.3 kDa), as has been observed previously (60).

**Supplementary figure 2.**
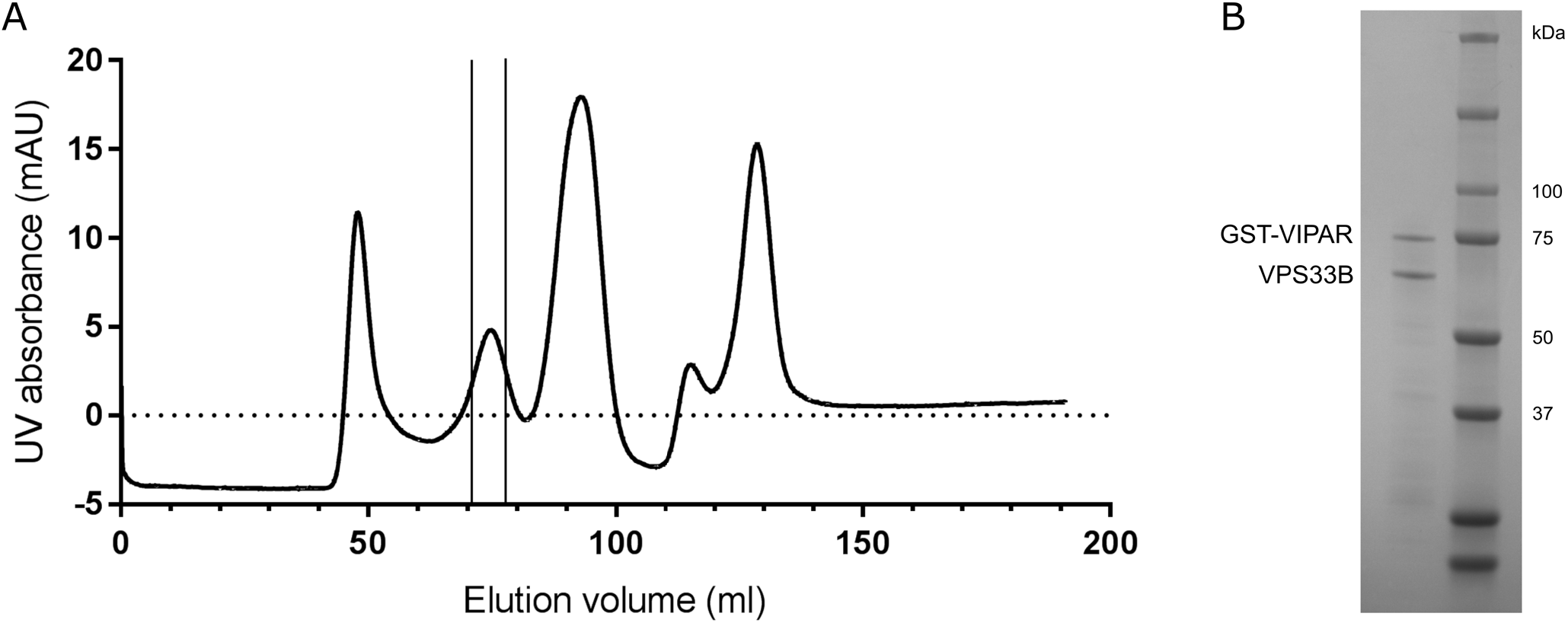
Co-expression and purification of VPS33B and GST-VIPAR. (A) Size-exclusion chromatogram (Superdex 200 10/300 GL) showing purification of the VPS33B/GST-VIPAR complex from *E. coli* following GSH affinity purification. Fractions from the indicated peak were pooled. (B) Pooled and concentrated fractions were analysed by SDS-PAGE. The indicated bands were confirmed by mass fingerprinting as indicated.

## Abbreviations

ARC: Arthrogryposis, Renal dysfunction, and Cholestasis syndrome
ARKID: Autosomal recessive Keratoderma-Ichthyosis-Deafness syndrome
CHEVI: class C Homologs in Endosome-Vesicle Interaction
CORVET: class C core vacuole/endosome tethering
EARP: endosome-associated recycling protein
GARP: Golgi associated retrograde protein
EGFP: GFP (enhanced) green fluorescent protein
HEK293T: human embryonic kidney 293T
HOPS: homotypic fusion and vacuole protein sorting
MS: mass spectrometry
PI3KC3: class III phosphatidylinositol 3-kinase
XLID: X-linked intellectual disability

## REFERENCES

1. D’Agostino, M., Risselada, H. J., Lürick, A., Ungermann, C., and Mayer, A. (2017) A tethering complex drives the terminal stage of SNARE-dependent membrane fusion. Nature 551, 634–638

2. Balderhaar, H. J. k., and Ungermann, C. (2013) CORVET and HOPS tethering complexes – coordinators of endosome and lysosome fusion. Journal of Cell Science 126, 1307–1316

3. Perini, E. D., Schaefer, R., Stöter, M., Kalaidzidis, Y., and Zerial, M. (2014) Mammalian CORVET is required for fusion and conversion of distinct early endosome subpopulations. Traffic 15, 1366–1389

4. van der Kant, R., Jonker, C. T., Wijdeven, R. H., Bakker, J., Janssen, L., Klumperman, J., and Neefjes, J. (2015) Characterization of the mammalian CORVET and HOPS complexes and their modular restructuring for endosome specificity. Journal of Biological Chemistry 290, 30280–30290

5. Klinger, C. M., Klute, M. J., and Dacks, J. B. (2013) Comparative genomic analysis of multi-subunit tethering complexes demonstrates an ancient pan-eukaryotic complement and sculpting in Apicomplexa. PloS one 8, e76278

6. Lachmann, J., Glaubke, E., Moore, P. S., and Ungermann, C. (2014) The Vps39-like TRAP1 is an effector of Rab5 and likely the missing Vps3 subunit of human CORVET. Cellular Logistics 4, e970840

7. van der Kant, R., Fish, A., Janssen, L., Janssen, H., Krom, S., Ho, N., Brummelkamp, T., Carette, J., Rocha, N., and Neefjes, J. (2013) Late endosomal transport and tethering are coupled processes controlled by RILP and the cholesterol sensor ORP1L. Journal of Cell Science 126, 3462–3474

8. Wartosch, L., Günesdogan, U., Graham, S. C., and Luzio, J. P. (2015) Recruitment of VPS33A to HOPS by VPS16 is required for lysosome fusion with endosomes and autophagosomes. Traffic 16, 727–742

9. Jiang, P., Nishimura, T., Sakamaki, Y., Itakura, E., Hatta, T., Natsume, T., and Mizushima, N. (2014) The HOPS complex mediates autophagosome-lysosome fusion through interaction with syntaxin 17. Molecular Biology of the Cell 25, 1327–1337

10. Gissen, P., Johnson, C. A., Gentle, D., Hurst, L. D., Doherty, A. J., O’kane, C. J., Kelly, D. A., and Maher, E. R. (2005) Comparative evolutionary analysis of VPS33 homologues: genetic and functional insights. Human Molecular Genetics 14, 1261–1270

11. Graham, S. C., Wartosch, L., Gray, S. R., Scourfield, E. J., Deane, J. E., Luzio, J. P., and Owen, D. J. (2013) Structural basis of Vps33A recruitment to the human HOPS complex by Vps16. Proceedings of the National Academy of Sciences of the USA 110, 13345–13350

12. Banushi, B., Forneris, F., Straatman-Iwanowska, A., Strange, A., Lyne, A.-M., Rogerson, C., Burden, J. J., Heywood, W. E., Hanley, J., Doykov, I., Straatman, K. R., Smith, H., Bem, D., Kriston-Vizi, J., Ariceta, G., Risteli, M., Wang, C., Ardill, R. E., Zaniew, M., Latka-Grot, J., Waddington, S. N., Howe, S. J., Ferraro, F., Gjinovci, A., Lawrence, S., Marsh, M., Girolami, M., Bozec, L., Mills, K., and Gissen, P. (2016) Regulation of post-Golgi LH3 trafficking is essential for collagen homeostasis. Nature Communications 7, 12111

13. Cullinane, A. R., Straatman-Iwanowska, A., Zaucker, A., Wakabayashi, Y., Bruce, C. K., Luo, G., Rahman, F., Gürakan, F., Utine, E., Özkan, T. B., Denecke, J., Vukovic, J., Di Rocco, M., Mandel, H., Cangul, H., Matthews, R. P., Thomas, S. G., Rappoport, J. Z., Arias, I. M., Wolburg, H., Knisely, A. S., Kelly, D. A., Müller, F., Maher, E. R., and Gissen, P. (2010) Mutations in VIPAR cause an arthrogryposis, renal dysfunction and cholestasis syndrome phenotype with defects in epithelial polarization. Nature Genetics 42, 303–312

14. Kondo, H., Maksimova, N., Otomo, T., Kato, H., Imai, A., Asano, Y., Kobayashi, K., Nojima, S., Nakaya, A., Hamada, Y., Irahara, K., Gurinova, E., Sukhomyasova, A., Nogovicina, A., Savvina, M., Yoshimori, T., Ozono, K., and Sakai, N. (2017) Mutation in VPS33A affects metabolism of glycosaminoglycans: a new type of mucopolysaccharidosis with severe systemic symptoms. Human Molecular Genetics 26, 173–183

15. Dursun, A., Yalnizoglu, D., Gerdan, O. F., Yucel-Yilmaz, D., Sagiroglu, M. S., Yuksel, B., Gucer, S., Sivri, S., and Ozgul, R. K. (2017) A probable new syndrome with the storage disease phenotype caused by the VPS33A gene mutation. Clinical dysmorphology 26, 1–12

16. Lo, B., Li, L., Gissen, P., Christensen, H., McKiernan, P. J., Ye, C., Abdelhaleem, M., Hayes, J. A., Williams, M. D., Chitayat, D., and Kahr, W. H. A. (2005) Requirement of VPS33B, a member of the Sec1/Munc18 protein family, in megakaryocyte and platelet α-granule biogenesis. Blood 106, 4159–4166

17. Urban, D., Li, L., Christensen, H., Pluthero, F. G., Chen, S. Z., Puhacz, M., Garg, P. M., Lanka, K. K., Cummings, J. J., Kramer, H., Wasmuth, J. D., Parkinson, J., and Kahr, W. H. A. (2012) The VPS33B-binding protein VPS16B is required in megakaryocyte and platelet α-granule biogenesis. Blood 120, 5032–5040

18. Gruber, R., Rogerson, C., Windpassinger, C., Banushi, B., Straatman-Iwanowska, A., Hanley, J., Forneris, F., Strohal, R., Ulz, P., and Crumrine, D. (2017) Autosomal Recessive Keratoderma-Ichthyosis-Deafness (ARKID) Syndrome Is Caused by VPS33B Mutations Affecting Rab Protein Interaction and Collagen Modification. Journal of Investigative Dermatology 137, 845–854

19. Spang, A. (2016) Membrane Tethering Complexes in the Endosomal System. Frontiers in Cell and Developmental Biology 4, 35

20. Lobingier, B. T., and Merz, A. J. (2012) Sec1/Munc18 protein Vps33 binds to SNARE domains and the quaternary SNARE complex. Molecular Biology of the Cell 23, 4611–4622

21. Zhang, Y. (2008) I-TASSER server for protein 3D structure prediction. BMC Bioinformatics 9, 40

22. Roux, K. J., Kim, D. I., Raida, M., and Burke, B. (2012) A promiscuous biotin ligase fusion protein identifies proximal and interacting proteins in mammalian cells. Journal of Cell Biology 196, 801–810

23. Teo, G., Liu, G., Zhang, J., Nesvizhskii, A. I., Gingras, A. C., and Choi, H. (2014) SAINTexpress: improvements and additional features in Significance Analysis of INTeractome software. Journal of proteomics 100, 37–43

24. Nickerson, D. P., Brett, C. L., and Merz, A. J. (2009) Vps-C complexes: gatekeepers of endolysosomal traffic. Current Opinion in Cell Biology 21, 543–551

25. Murray, J. T., Panaretou, C., Stenmark, H., Miaczynska, M., and Backer, J. M. (2002) Role of Rab5 in the Recruitment of hVps34/p150 to the Early Endosome. Traffic 3, 416–427

26. Stein, M.-P., Feng, Y., Cooper, K. L., Welford, A. M., and Wandinger-Ness, A. (2003) Human VPS34 and p150 are Rab7 Interacting Partners. Traffic 4, 754–771

27. Backer, J. M. (2016) The intricate regulation and complex functions of the Class III phosphoinositide 3-kinase Vps34. Biochemical Journal 473, 2251–2271

28. Lu, J., He, L., Behrends, C., Araki, M., Araki, K., Jun Wang, Q., Catanzaro, J. M., Friedman, S. L., Zong, W.-X., Fiel, M. I., Li, M., and Yue, Z. (2014) NRBF2 regulates autophagy and prevents liver injury by modulating Atg14L-linked phosphatidylinositol-3 kinase III activity. Nature Communications 5, 3920

29. Cao, Y., Wang, Y., Abi Saab, Widian F., Yang, F., Pessin, Jeffrey E., and Backer, Jonathan M. (2014) NRBF2 regulates macroautophagy as a component of Vps34 Complex I. Biochemical Journal 461, 315–322

30. Zhong, Y., Morris, D. H., Jin, L., Patel, M. S., Karunakaran, S. K., Fu, Y.-J., Matuszak, E. A., Weiss, H. L., Chait, B. T., and Wang, Q. J. (2014) Nrbf2 protein suppresses autophagy by modulating Atg14L protein-containing Beclin 1-Vps34 complex architecture and reducing intracellular phosphatidylinositol-3 phosphate levels. Journal of Biological Chemistry 289, 26021–26037

31. Liang, C., Lee, J.-s., Inn, K.-S., Gack, M. U., Li, Q., Roberts, E. A., Vergne, I., Deretic, V., Feng, P., and Akazawa, C. (2008) Beclin1-binding UVRAG targets the class C Vps complex to coordinate autophagosome maturation and endocytic trafficking. Nature Cell Biology 10, 776–787

32. Sun, Q., Westphal, W., Wong, K. N., Tan, I., and Zhong, Q. (2010) Rubicon controls endosome maturation as a Rab7 effector. Proceedings of the National Academy of Sciences of the USA 107, 19338–19343

33. Sun, Q., Zhang, J., Fan, W., Wong, K. N., Ding, X., Chen, S., and Zhong, Q. (2011) The RUN domain of rubicon is important for hVps34 binding, lipid kinase inhibition, and autophagy suppression. Journal of Biological Chemistry 286, 185–191

34. Cebrian, I., Visentin, G., Blanchard, N., Jouve, M., Bobard, A., Moita, C., Enninga, J., Moita, Luis F., Amigorena, S., and Savina, A. (2011) Sec22b Regulates Phagosomal Maturation and Antigen Crosspresentation by Dendritic Cells. Cell 147, 1355–1368

35. Petkovic, M., Jemaiel, A., Daste, F., Specht, C. G., Izeddin, I., Vorkel, D., Verbavatz, J. M., Darzacq, X., Triller, A., Pfenninger, K. H., Tareste, D., Jackson, C. L., and Galli, T. (2014) The SNARE Sec22b has a non-fusogenic function in plasma membrane expansion. Nature Cell Biology 16, 434–444

36. Dai, J., Lu, Y., Wang, C., Chen, X., Fan, X., Gu, H., Wu, X., Wang, K., Gartner, T. K., Zheng, J., Chen, G., Wang, X., and Liu, J. (2016) Vps33b regulates Vwf-positive vesicular trafficking in megakaryocytes. The Journal of pathology 240, 108–119

37. Phillips-Krawczak, C. A., Singla, A., Starokadomskyy, P., Deng, Z., Osborne, D. G., Li, H., Dick, C. J., Gomez, T. S., Koenecke, M., Zhang, J.-S., Dai, H., Sifuentes-Dominguez, L. F., Geng, L. N., Kaufmann, S.H., Hein, M. Y., Wallis, M., McGaughran, J., Gecz, J., Sluis, B. v. d., Billadeau, D. D., and Burstein, E. (2015) COMMD1 is linked to the WASH complex and regulates endosomal trafficking of the copper transporter ATP7A. Molecular Biology of the Cell 26, 91–103

38. Wan, C., Borgeson, B., Phanse, S., Tu, F., Drew, K., Clark, G., Xiong, X., Kagan, O., Kwan, J., Bezginov, A., Chessman, K., Pal, S., Cromar, G., Papoulas, O., Ni, Z., Boutz, D. R., Stoilova, S., Havugimana, P. C., Guo, X., Malty, R. H., Sarov, M., Greenblatt, J., Babu, M., Derry, W. B., R. Tillier, E., Wallingford, J. B., Parkinson, J., Marcotte, E. M., and Emili, A. (2015) Panorama of ancient metazoan macromolecular complexes. Nature 525, 339–344

39. Bartuzi, P., Billadeau, D. D., Favier, R., Rong, S., Dekker, D., Fedoseienko, A., Fieten, H., Wijers, M., Levels, J. H., Huijkman, N., Kloosterhuis, N., van der Molen, H., Brufau, G., Groen, A. K., Elliott, A. M., Kuivenhoven, J. A., Plecko, B., Grangl, G., McGaughran, J., Horton, J. D., Burstein, E., Hofker, M. H., and van de Sluis, B. (2016) CCC-and WASH-mediated endosomal sorting of LDLR is required for normal clearance of circulating LDL. Nature Communications 7, 10961

40. Harbour, Michael E., Breusegem, Sophia Y., and Seaman, Matthew N. J. (2012) Recruitment of the endosomal WASH complex is mediated by the extended ‘tail’ of Fam21 binding to the retromer protein Vps35. Biochemical Journal 442, 209–220

41. McNally, K. E., Faulkner, R., Steinberg, F., Gallon, M., Ghai, R., Pim, D., Langton, P., Pearson, N., Danson, C. M., Nägele, H., Morris, L. L., Singla, A., Overlee, Brittany L., Heesom, K. J., Sessions, R., Banks, L., Collins, B. M., Berger, I., Billadeau, D. D., Burstein, E., and Cullen, P. J. (2017) Retriever is a multiprotein complex for retromer-independent endosomal cargo recycling. Nature Cell Biology 19, 1214–1225

42. Schindler, C., Chen, Y., Pu, J., Guo, X., and Bonifacino, J. S. (2015) EARP is a multisubunit tethering complex involved in endocytic recycling. Nature Cell Biology 17, 639–650

43. Voineagu, I., Huang, L., Winden, K., Lazaro, M., Haan, E., Nelson, J., McGaughran, J., Nguyen, L. S., Friend, K., Hackett, A., Field, M., Gecz, J., and Geschwind, D. (2011) CCDC22: a novel candidate gene for syndromic X-linked intellectual disability. Molecular Psychiatry 17, 4–7

44. Starokadomskyy, P., Gluck, N., Li, H., Chen, B., Wallis, M., Maine, G. N., Mao, X., Zaidi, I. W., Hein, M. Y., McDonald, F. J., Lenzner, S., Zecha, A., Ropers, H.-H., Kuss, A. W., McGaughran, J., Gecz, J., and Burstein, E. (2013) CCDC22 deficiency in humans blunts activation of proinflammatory NF-κB signaling. The Journal of Clinical Investigation 123, 2244–2256

45. Bonifacino, J. S., and Hierro, A. (2011) Transport according to GARP: receiving retrograde cargo at the trans-Golgi network. Trends in Cell Biology 21, 159–167

46. Huttlin, E. L., Bruckner, R. J., Paulo, J. A., Cannon, J. R., Ting, L., Baltier, K., Colby, G., Gebreab, F., Gygi, M. P., Parzen, H., Szpyt, J., Tam, S., Zarraga, G., Pontano-Vaites, L., Swarup, S., White, A. E., Schweppe, D. K., Rad, R., Erickson, B. K., Obar, R. A., Guruharsha, K. G., Li, K., Artavanis-Tsakonas, S., Gygi, S. P., and Harper, J. W. (2017) Architecture of the human interactome defines protein communities and disease networks. Nature 545, 505–509

47. Bach, H., Papavinasasundaram, K. G., Wong, D., Hmama, Z., and Av-Gay, Y. (2008) Mycobacterium tuberculosis virulence is mediated by PtpA dephosphorylation of human vacuolar protein sorting 33B. Cell Host & Microbe 3, 316–322

48. Akbar, M. A., Tracy, C., Kahr, W. H., and Krämer, H. (2011) The full-of-bacteria gene is required for phagosome maturation during immune defense in Drosophila. Journal of Cell Biology 192, 383–390

49. Akbar, M. A., Mandraju, R., Tracy, C., Hu, W., Pasare, C., and Krämer, H. (2016) ARC Syndrome-Linked Vps33B Protein Is Required for Inflammatory Endosomal Maturation and Signal Termination. Immunity 45, 267–279

50. Couzens, A. L., Knight, J. D., Kean, M. J., Teo, G., Weiss, A., Dunham, W. H., Lin, Z. Y., Bagshaw, R. D., Sicheri, F., Pawson, T., Wrana, J. L., Choi, H., and Gingras, A. C. (2013) Protein interaction network of the mammalian Hippo pathway reveals mechanisms of kinase-phosphatase interactions. Science signaling 6, rs15

51. Teo, H., Perisic, O., González, B., and Williams, R. L. (2004) ESCRT-II, an Endosome-Associated Complex Required for Protein Sorting: Crystal Structure and Interactions with ESCRT-III and Membranes. Developmental Cell 7, 559–569

52. Chapat, C., Jafarnejad, S. M., Matta-Camacho, E., Hesketh, G. G., Gelbart, I. A., Attig, J., Gkogkas, C. G., Alain, T., Stern-Ginossar, N., Fabian, M. R., Gingras, A. C., Duchaine, T. F., and Sonenberg, N. (2017) Cap-binding protein 4EHP effects translation silencing by microRNAs. Proceedings of the National Academy of Sciences of the USA 114, 5425–5430

53. Liu, G., Zhang, J., Larsen, B., Stark, C., Breitkreutz, A., Lin, Z. Y., Breitkreutz, B. J., Ding, Y., Colwill, K., Pasculescu, A., Pawson, T., Wrana, J. L., Nesvizhskii, A. I., Raught, B., Tyers, M., and Gingras, A. C. (2010) ProHits: integrated software for mass spectrometry-based interaction proteomics. Nature biotechnology 28, 1015–1017

54. Liu, G., Knight, J. D., Zhang, J. P., Tsou, C. C., Wang, J., Lambert, J. P., Larsen, B., Tyers, M., Raught, B., Bandeira, N., Nesvizhskii, A. I., Choi, H., and Gingras, A. C. (2016) Data Independent Acquisition analysis in ProHits 4.0. Journal of proteomics 149, 64–68

55. Knight, J. D. R., Choi, H., Gupta, G. D., Pelletier, L., Raught, B., Nesvizhskii, A. I., and Gingras, A. C. (2017) ProHits-viz: a suite of web tools for visualizing interaction proteomics data. Nature methods 14, 645–646

56. Deutsch, E. W., Csordas, A., Sun, Z., Jarnuczak, A., Perez-Riverol, Y., Ternent, T., Campbell, D. S., Bernal-Llinares, M., Okuda, S., Kawano, S., Moritz, R. L., Carver, J. J., Wang, M., Ishihama, Y., Bandeira, N., Hermjakob, H., and Vizcaíno, J. A. (2017) The ProteomeXchange consortium in 2017: supporting the cultural change in proteomics public data deposition. Nucleic Acids Research 45, D1100–D1106

57. Hunter, M. R., Scourfield, E. J., Emmott, E., and Graham, S. C. (2017) VPS18 recruits VPS41 to the human HOPS complex via a RING–RING interaction. Biochemical Journal 474, 3615–3626

58. Zhu, G.-d., Salazar, G., Zlatic, S. A., Fiza, B., Doucette, M. M., Heilman, C. J., Levey, A. I., Faundez, V., and L’Hernault, S. W. (2009) SPE-39 Family Proteins Interact with the HOPS Complex and Function in Lysosomal Delivery. Molecular Biology of the Cell 20, 1223–1240

59. Schou, K. B., Andersen, J. S., and Pedersen, L. B. (2014) A divergent calponin homology (NN–CH) domain defines a novel family: implications for evolution of ciliary IFT complex B proteins. Bioinformatics 30, 899–902

60. Montalbano, J., Jin, W., Sheikh, M. S., and Huang, Y. (2007) RBEL1 Is a Novel Gene That Encodes a Nucleocytoplasmic Ras Superfamily GTP-binding Protein and Is Overexpressed in Breast Cancer. Journal of Biological Chemistry 282, 37640–37649

